# Diurnal variation of circulating interleukin-6 in humans: a meta-analysis

**DOI:** 10.1101/042507

**Authors:** Gustav Nilsonne, Mats Lekander, Torbjörn Åkerstedt, John Axelsson, Michael Ingre

**Affiliations:** Stockholm University, Stress Research Institute, Stockholm, Sweden; Karolinska Institutet, Department of Clinical Neuroscience, Stockholm, Sweden

## Abstract

The pleiotropic cytokine interleukin-6 (IL-6) has been proposed to contribute to circadian regulation of sleepiness by increasing in the blood at night to signal for sleepiness. Earlier studies have reported diurnal variations of IL-6, but phase estimates are conflicting. We have therefore performed a meta-analysis on the diurnal variation of circulating IL-6. Studies were included if they reported circulating levels of IL-6 recorded at least twice within 24 hours in the same individual. A systematic search resulted in the inclusion of 43 studies with 56 datasets, for a total of 1100 participants. Individual participant data were available from 4 datasets with a total of 56 participants. Mixed-effects meta-regression modelling confirmed that IL-6 varied across the day, the most conspicuous effect being a trough in the morning. These results stand in contrast to earlier findings of a peak in the evening or night, and suggest that diurnal variation should be taken into account in order to avoid confounding in studies of IL-6 in plasma or serum.

## Introduction

Sleepiness is regulated in humans by two main processes: the circadian process, which makes us sleepier in the night, and the homeostatic process, which causes sleepiness to increase with time awake [1]. It has been proposed that interleukin-6, a pleiotropic cytokine, participates in circadian sleepiness regulation by increasing at night in the blood and inducing sleepiness through signalling in the brain [2–6]. Early studies of diurnal variation of IL-6 in humans found a peak in the night-time [7,8], and it is this observational relationship that forms the main line of evidence for a regulatory effect of circulating IL-6 on sleepiness. However, further studies have since found peaks at different times of the day or have found no peaks at all. Fig 1 shows locations of peaks and troughs that have been estimated in the literature so far. Notably, estimates have ranged quite widely. Nonetheless, the general impression of these earlier claims is consistent with an increase of IL-6 levels in the night-time.

**Fig 1.**
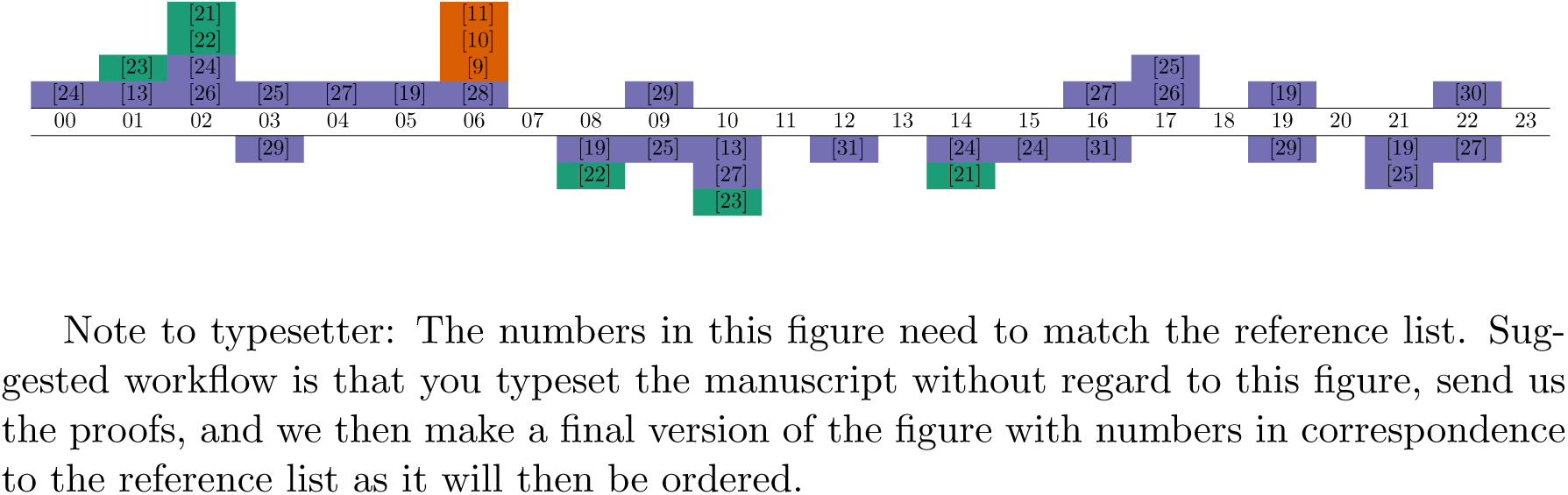
Estimates of phase reported in earlier literature. Every count represents one claim of having located a peak (box above time-line) or a trough (box below time-line) in a dataset. Blue: Studies included in quantitative review. Green: Studies not included in quantitative review. Orange: Meta-analysis. Review papers are not included.

One previous meta-analysis of IL-6 and time of day has been reported [9] (and published again in [10] and [11]). This meta-analysis focused on the diurnal variation of interleukin-6 in rheumatoid arthritis, but also included an estimate for healthy control participants from 11 studies. Data inclusion procedures were informal; no systematic method for identifying and including data was reported. The main finding in healthy participants was an increase if IL-6 from the evening, continuing during the night, followed by a drop in the morning. The pattern in patients with rheumatoid arthritis was similar, but with a more pronounced peak in the early morning before levels started to fall.

Thus, the observational relationship between IL-6 and time of day in healthy humans has important implications for the theoretical understanding of immune-brain interactions in sleepiness regulation, but there is no consensus on estimates of phase. Therefore, we have performed a meta-analysis, aiming to investigate the diurnal variation of IL-6 in the blood.

## Materials and Methods

### Literature search and data acquisition

The PubMed database was searched using the terms “interleukin-6 AND (sleep OR diurnal OR circadian)”, and the limit “human”. The search was last updated on 2016-01-03. Records were reviewed by one investigator (GN). Studies were included if they reported IL-6 in plasma or serum from healthy participants with a time-course including two or more time-points within 24 hours. Fig 2 shows a flowchart of data inclusion. Table 1 shows characteristics of included studies. Table 2 lists studies that fulfulled inclusion criteria but which could nonetheless not be included. The most common reason was that data could not be estimated (*k* = 25). Of these 25 studies, 7 reported that data were largely or entirely under the assay detection limit. In the remaining cases, data could not be estimated because they were given as a difference score (*k* = 5), because they were not shown (*k* = 4), because time of day was not given (*k* = 3), or for other reasons, specified in table 2 (*k* = 6). Additionally, seven studies were excluded due to duplicate publication of data, and four studies were excluded because the reported levels of IL-6 were very high and therefore judged not to represent levels consistent with physiological regulation or variation in healthy humans. Of these studies, one reported one participant, whose IL-6 levels increased ten-fold after venous catheterization [72], and we judged that this change was not representative of diurnal variation. Another study reported ten participants with mean plasma IL-6 levels of about 10-30 pg/ml over the course of two days [73]. This is about ten times higher than expected for healthy participants, raising questions about the validity of the absolute values. We judged that these measures may not accurately reflect the target outcome, and that they would unduly influence the regression model on account of the very high values. They were therefore excluded. Finally, two studies [83,84] reported 40 and 60 participants, possibly with overlapping samples, both with mean values of about 35 and 55 pg/ml at 09:00 and 02:00, respectively. Because these levels were so much higher than expected for healthy participants, these studies were also excluded. Thirteen studies were included from sources other than the PubMed results. These were found mostly because they were cited by papers identified in the literature search, and in some cases because they were known to us on beforehand.

**Fig 2.**
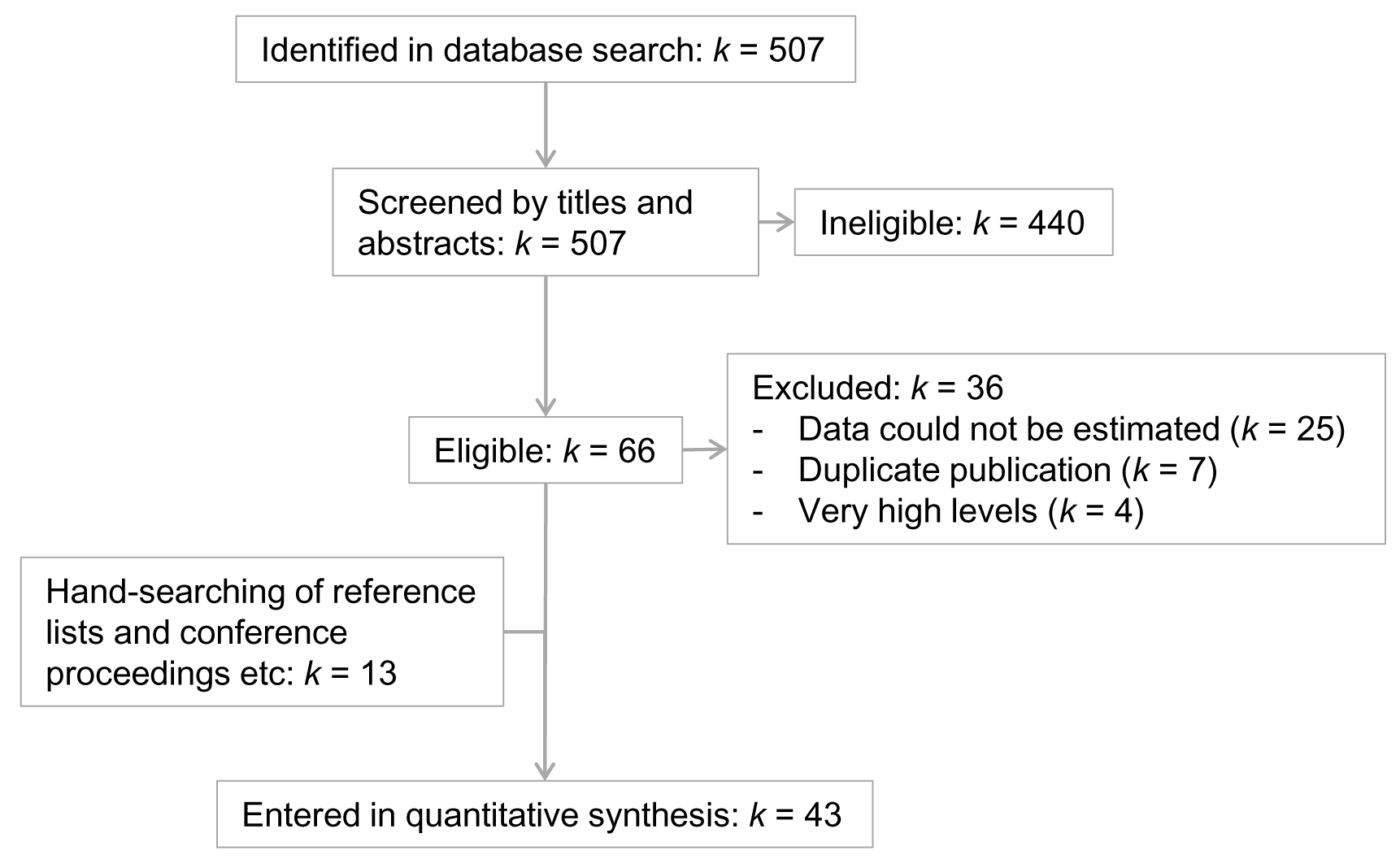
Data inclusion. Some of the 43 included studies contained more than one dataset. The final number of datasets was 56 (see table 1).

**Table 1.** Characteristics of included studies. Where studies reported several datasets, these are specified separately.

| 1<sup>st</sup> Author | Year | Ref. | $n_{subjects}$ | $n_{timepoints}$ | Regression weight, % | Notes |
| --- | --- | --- | --- | --- | --- | --- |
| Sothorn | 1995 | [13] | 11 | 8 | 0.99 | a |
| Späth-Schwalbe | 1998 | [32] | 16 | 4 | 1.01 |  |
| Dugué | 1998 | [33] | 22 | 3 | 1.21 |  |
| Vgontzas | 1999 | [19] | 8 | 24 | 1.24 |  |
| Ündar | 1999 | [28] | 10 | 7 | 0.84 |  |
| Späth-Schwalbe | 2000 | [34] | 18 | 3 | 0.99 |  |
| Redwine | 2000 | [35] | 31 | 15 | 3.80 | b |
| Haack | 2000 | [36] | 20 | 9 | 1.90 |  |
| Vgontzas | 2000 | [37] | 12; 11 | 2; 2 | 0.54; 0.49 | c |
| Baker | 2001 | [38] | 8 | 6 | 0.62 |  |
| Haack | 2001 | [39] | 10 | 10 | 1.00 |  |
| Haack | 2002 | [16] | 12 | 25 | 1.99 |  |
| Vgontzas | 2002 | [26] | 11 | 24 | 1.71 |  |
| Vgontzas | 2003 | [25] | 15; 13 | 48; 48 | 3.29; 2.85 |  |
| Alberti | 2003 | [40] | 20 | 2 | 0.49 |  |
| Irwin | 2004 | [41] | 15 | 4 | 0.95 |  |
| Vgontzas | 2004 | [42] | 25 | 48 | 5.49 |  |
| Szczudlik | 2004 | [43] | 17 | 4 | 1.08 |  |
| Alesci | 2005 | [24] | 9 | 22 | 1.34 |  |
| Mehra | 2006 | [44] | 150; 23; 12; 35; 30 | 2; 2; 2; 2; 2 | 6.72; 1.03; 0.54; 1.56; 1.34 | d |
| Burgos | 2006 | [29] | 11 | 8 | 0.99 |  |
| Vgontzas | 2007 | [45] | 20; 20 | 48; 48 | 4.39; 4.39 | e |
| Frey | 2007 | [46] | 19 | 15 | 2.33 | b |
| Lindahl | 2007 | [47] | 14 | 4 | 0.89 |  |
| Miles | 2008 | [31] | 51 | 5 | 3.61 |  |
| Vgontzas | 2008 | [48] | 15; 13 | 48; 48 | 3.29; 2.85 |  |
| Knudsen | 2008 | [30] | 15 | 8 | 1.14 | a, f |
| Benedict | 2009 | [49] | 17 | 15 | 2.09 |  |
| Peeling | 2009 | [50] | 8 | 5 | 0.57 |  |
| Phillips | 2010 | [51] | 7 | 3; 3 | 0.38; 0.38 | g |
| Rief | 2010 | [52] | 60; 52 | 2; 2 | 2.69; 2.33 |  |
| Gill | 2010 | [53] | 14 | 13 | 1.60 |  |
| Chennaoui | 2011 | [54] | 12 | 6 | 0.93 |  |
| Pledge | 2011 | [55] | 6 | 4 | 0.76 |  |
| Crispim | 2012 | [56] | 6; 7; 9 | 7; 7; 7 | 0.50; 0.59; 0.75 |  |
| Voderholzer | 2012 | [57] | 16 | 6 | 1.24 |  |
| Grigoleit | 2012 | [58] | 10 | 7 | 0.84 |  |
| Abdelmalek | 2013 | [59] | 12 | 2 | 0.54 |  |
| Pejovic | 2013 | [60] | 30 | 24 | 4.65 |  |
| Lekander | 2013 | [61] | 9 | 40 | 1.30 | a |
| Agorastos | 2014 | [27] | 11 | 24 | 1.71 |  |
| Kritikou | 2014 | [62] | 18; 21 | 24; 24 | 2.79; 3.26 |  |
| Karshikoff | 2015 | [14] | 21 | 4 | 1.31 | a, h |
| Total |  |  | 1100 | 789 | 100 |  |

a: Individual participant data were available. b: Some data were given in time relative to sleep onset or wake-up, and were re-coded using mean chronological time as a best approximation. c: Averaged over 3 consecutive days. d: 358 of 385 participants were included in analyses of IL-6. Final $n$ for each sub-group was not given, and was therefore conservatively coded as the lowest possible $n$ in each sub-group. Error bars were denoted as standard deviation, but were coded as standard errors because they were incredibly small for standard deviations. e: Each dataset was said to have 50% of the total participants ( $n = 41$ ), and both were conservatively coded as $n = 20$ . f: 15 of 16 participants could be identified in the graph. g: The same 7 participants were included twice with a 10-week interval, yielding two different data sets. h: We have previously published data from this study [63].

**Table 2.** Characteristics of excluded studies.

| 1<sup>st</sup> author | Year | Ref. | $n_{subjects}$ | Reason for exclusion |
| --- | --- | --- | --- | --- |
| Lemmer | 1992 | [64] | 12 | Data could not be estimated (below detection limit) |
| Gudewill | 1992 | [7] | 12 | Data could not be estimated (given as counts) |
| Pollmächer | 1993 | [65] | 15 | Data could not be estimated (not shown; were “close to assay detection limit”) |
| Bauer | 1994 | [8] | 5 | Data could not be estimated (given as arbitrary units) |
| Dinges | 1994 | [66] | 20 | Data could not be estimated (not shown) |
| Arvidson | 1994 | [67] | 10 | Data could not be estimated (below detection limit) |
| Seiler | 1994 | [15] | 6 | Data could not be estimated (clock time not given, and too low resolution) |
| Seiler | 1995 | [68] | 6 | Data same as in [15] |
| Sothorn | 1995 | [23] | 10 | Data same as in [13] |
| Pollmächer | 1996 | [69] | 20 | Data could not be estimated (given as difference between treatments) |
| Korth | 1996 | [70] | 20 | Data could not be estimated (too low resolution) |
| Crofford | 1997 | [71] | 5 | Data could not be estimated (largely below detection limit) |
| Gudmundsson | 1997 | [72] | 1 | Very high levels |
| Born | 1997 | [73] | 10 | Very high levels |
| Lissoni | 1998 | [74] | 10 | Data could not be estimated (largely below detection limit) |
| Bornstein | 1998 | [75] | 9 | Data could not be estimated (not shown) |
| Hermann | 1998 | [76] | 10 | Data could not be estimated (too low resolution) |
| Kanabrocki | 1999 | [21] | 11 | Data same as in [13] |
| Genesca | 2000 | [77] | 8 | Data could not be estimated (largely below detection limit) |
| Mastorakos | 2000 | [78] | 5 | Data same as in [71] |
| Mullington | 2000 | [79] | 19 | Data could not be estimated (given as change between conditions), also possibly same as in [70] |
| Johansson | 2000 | [80] | 18 | Data could not be estimated (largely below detection limit) |
| Shaw | 2001 | [81] | 10 | Data could not be estimated (largely below detection limit) |
| Lange | 2002 | [82] | 18 | Data could not be estimated (given as z-transformed change between time points) |
| Domínguez-Rodríguez | 2003 | [83] | 40 | Very high levels |
| Domínguez-Rodríguez | 2004 | [84] | 60 | Very high levels and possibly overlapping with [83] |
| Haack | 2007 | [85] | 18 | Data could not be estimated (given as change between conditions) |
| Eisenberger | 2009 | [86] | 16 | Data could not be estimated (too low resolution) |
| Eisenberger | 2010 | [87] | 16 | Data same as in [86] |
| Haimovich | 2010 | [88] | 2 | Data could not be estimated (given as change between conditions) |
| Sauvet | 2010 | [89] | 12 | Data same as in [54] |
| Grigoleit | 2011 | [90] | 34 | Data could not be estimated (control condition not shown) |
| Miles | 2012 | [91] | 30 | Data same as in [31] |
| Schrepf | 2014 | [92] | 28 | Data could not be estimated (time of sampling not shown) |
| Wegner | 2014 | [93] | 18 | Data could not be estimated (too low resolution) |
| Scott | 2015 | [94] | 14 | Data could not be estimated (clock time not given) |
| Total |  |  | 468 |  |

The total number of participants does not count twice those participants included in duplicate reports.

Data were estimated from published tables or from graphs using GetData Graph Digitizer, version 2.25.0.25 (getdata-graph-digitizer.com). Error bars were assumed to represent standard errors unless otherwise indicated. Time of day was coded, as well as sleep or wake, time asleep, and time awake. For studies reporting participants kept in the lab overnight, data obtained at the time-point when the lights-out period began were coded as awake and data from the time-point when the lights-out period ended were coded as asleep. When times for falling asleep and waking up were not recorded or not reported, we assumed that they were 23:00 and 07:00. When applicable, time from catheter insertion was also coded. Unless otherwise specified, serial sampling with more than two samples within the same 24-hour period was assumed to have been performed using an indwelling catheter inserted at the first sampling time point. Data recorded during sleep deprivation were not included. IL-6 data were ln-transformed to better approximate a normal distribution. For datasets where individual participant data were not available, transformation was performed as described in [12].

Individual participant data were available from 4 datasets, which were coded separately (see table 1). In these datasets, data points below assay detection limits (meaning lowest known point of assay linear range) were conservatively re-coded to the value of the detection limit. In Sothern 1995 [13], 7 values out of 88 (8%), ranging from 0.5 to 0.96 pg/ml, were re-coded to 1 pg/ml. In Karshikoff 2015 [14], 23 values out of 83 (28%), ranging from 0.01 to 0.88 pg/ml, were re-coded to 0.9 pg/ml.

Ethical approval was not required. The study protocol was not registered. All data and the full analysis code are freely available at https://github.com/GNilsonne/IL6_diurnal.

### Meta-analysis

To investigate the diurnal time course of IL-6 in plasma and possible moderator variables, we used hierarchical mixed-effects models. This approach allows for more complex model fitting and is expected to have higher statistical power compared to fitting models separately in each dataset and then analysing summary measures such as acrophase and amplitude. Diurnal variation was investigated by fitting cosinor functions with periods of 24, 12, and 6 hours. Time from catheterisation was included with a random slope for each data set in order to account for the proposed effect that catheterization induces higher values in blood drawn from the catheter [15,16]. Effects of sleep were investigated exploratively with a binary factor for sleep/wake, as well as time asleep and time awake. Datasets were weighted by the number of participants multiplied by the square root of the number of time-points in each study. Models were compared using likelihood ratio tests. Analyses were performed using R version 3.2.0 [17] with the nlme package [18].

## Results

### Diurnal variation of IL-6

First, we fitted a null model including only time from catheterization and a random intercept for each dataset. Fig 3 shows residuals after these effects have been accounted for, suggesting that there remains variation to explain. The distribution of these residuals suggests a morning trough in IL-6 levels (Fig 3). Next, we compared a model with a 24 h cosinor function to the null model. The 24 h cosinor model fit better (log likelihood −483.5 vs −528.8, *p* < 0.0001, Fig 4). We then added another cosinor function with 12 h period. We did this for two reasons. The first reason was that addition of shorter periods allows a better estimation of non-sinusoidal effects, albeit at a cost of higher risk of overfitting. The second reason was that 12 h periods have been proposed by earlier investigators [19], and we considered that these claims should be tested. The model with both 24 h and 12 h cosinor functions fit better than the model with only the 24 h period (log likelihood −460.1 vs −483.5, *p* < 0.0001, Fig 4). Next, we exploratively investigated the addition of yet another cosinor with a 6 h period, but that did not improve model fit (log likelihood −457.7 vs −460.1, *p* = 0.09, prediction not shown).

**Fig 3.**
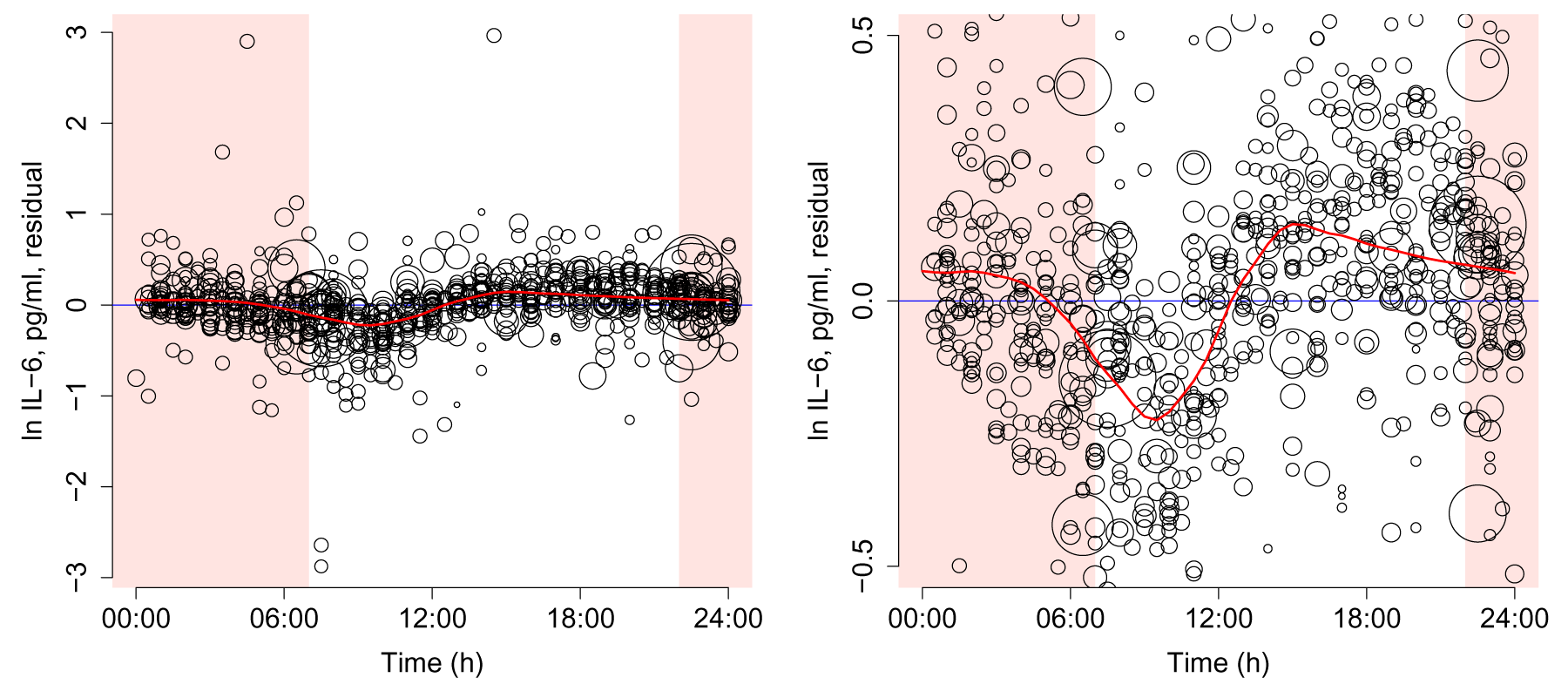
Residuals from null model. Data points sized by regression weight. In the null model, a random intercept for each study and a linear effect of time from catheterization have been included. Therefore, these residuals show the putative diurnal variation to be modeled. To explore this variation, we fitted a weighted LOESS curve (red line). This curve shows a trough in the morning. Note that the shape of the LOESS curve depends on the smoothing parameter. It is therefore possible to generate different LOESS curves from the same data, and not all of them show a peak in the early afternoon. The LOESS curve was fitted on three repeated days of the same data, and the curve for the second day shown, to ensure that the estimates would meet at 00:00 and 24:00. Time from 22:00 to 07:00 is shaded to indicate the night. Left: All data points shown. Right: Y axis range restricted to increase resolution.

**Fig 4.**
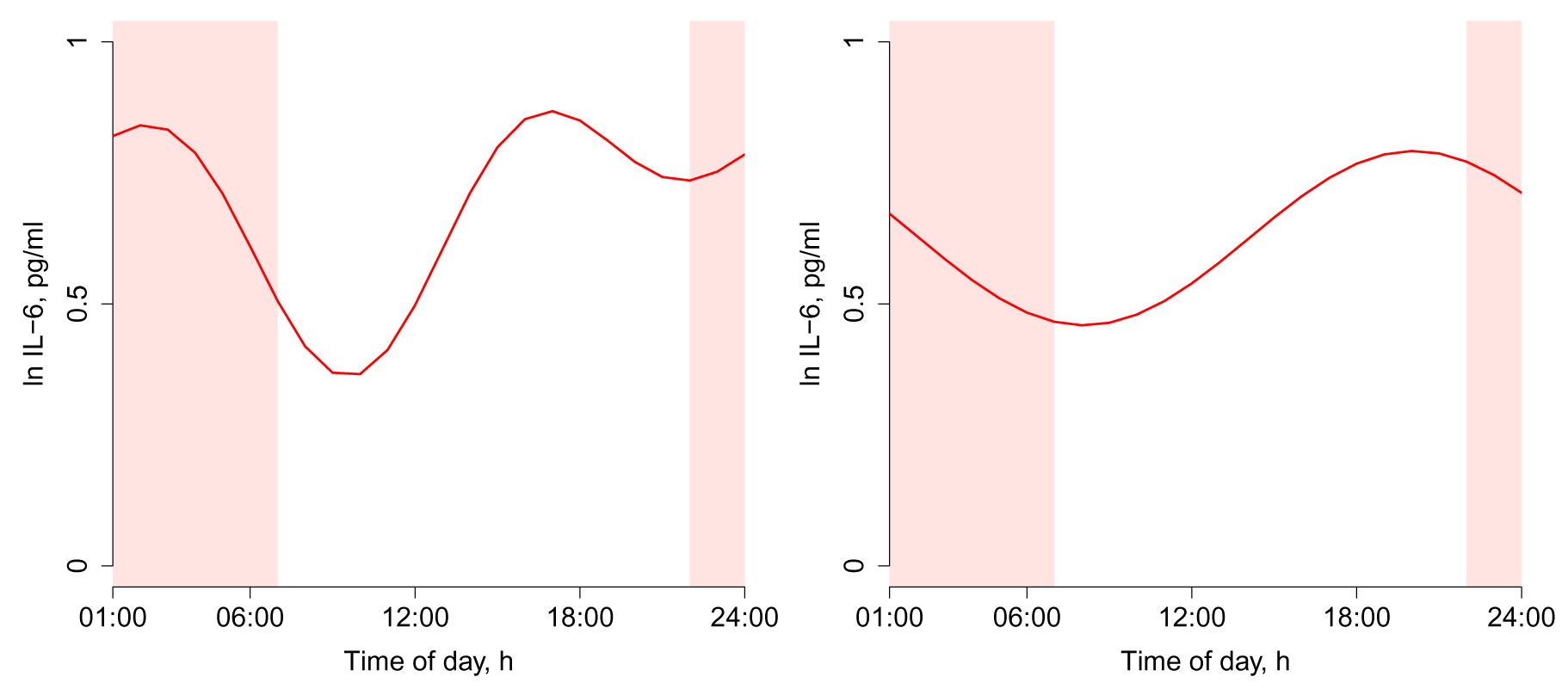
Predicted diurnal time courses from meta-regression models. Left: Best-fitting model including cosinor functions with 24 and 12 h periods. Right: Model including only 24 h period. Time from 22:00 to 07:00 is shaded to indicate the night.

Attempting to disentangle diurnal variation from effects of sleep, we investigated the addition of model effects for sleep (asleep/awake), time asleep (hours since sleep onset), and time awake (hours since wake onset). Starting with the best-fitting model including 24 and 12 h periods, we found that the addition of sleep, time asleep and time awake, or all three variables, did not improve model fit (log likelihoods −459.9, −459.2, and −459.0, respectively, vs −460.1, with *p* values 0.80, 0.36, and 0.36). Finally, we investigated the addition of sleep, time asleep and time awake, and all three variables, to the model with 24 h period. When comparing these models to the best-fitting model with 24 and 12 h periods, we saw worse fit (log likelihoods −475.5, −471.5, and −471.5, vs −460.1, with *p* values ≤ 0.0001). When comparing to the model with 24 h period only, we found that the addition of sleep, time asleep and time awake, or all three variables, yielded better-fitting models (log likelihoods −475.5, −471.5, and −471.5, vs −483.5, *p* values 0. 0003, < 0.0001, and < 0.0001).

The best-fitting model with 24 and 12 h periods had a conspicuous trough between 09:00 and 10:00 in the morning and a second less pronounced trough close to 22:00, and two peaks located close to 17:00 and 02:00 (Fig 4). Since diurnal rhythms are commonly investigated using cosinor functions with 24 h periods, we show predictions from this simpler model too (Fig 4). This model estimated bathyphase (lowest point) at 08:05 and acrophase (highest point) at 20:05, with an amplitude of 0.166. All the included datasets, with predictions from the best-fitting model, are shown in figures 5 and 6. Individual participant data are shown in figure 7, for those four studies from which individual participant data were available.

**Fig 5.**
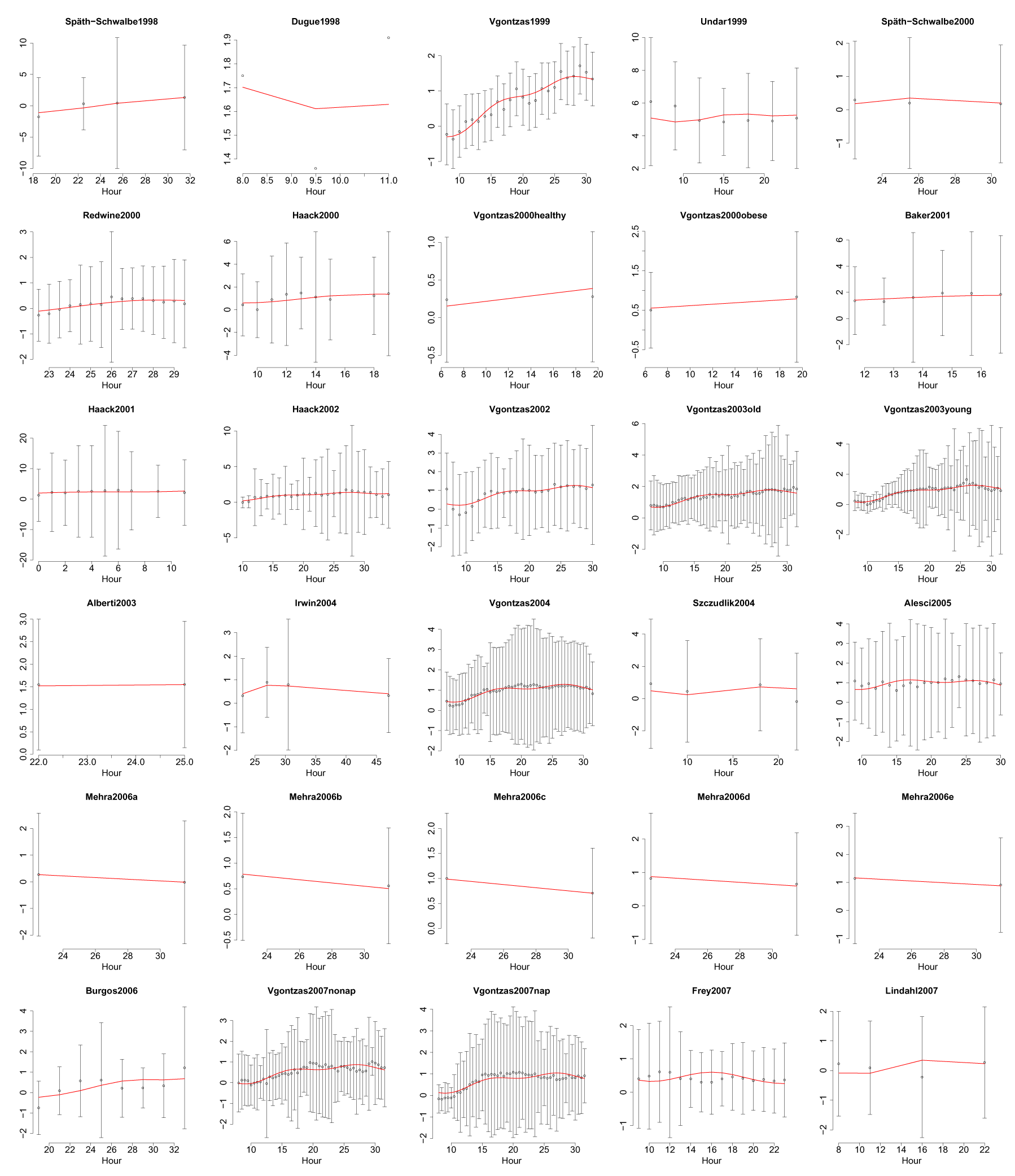
Data and fitted time courses, showing the first 30 out of 56 datasets. Data are shown as estimated from original publications, with error bars showing standard deviations. Y axes show ln IL-6 (pg/ml) throughout. Hours are in chronological time where 1 is 01:00 on the first day. Red lines show predictions from the best-fitting model.

**Fig 6.**
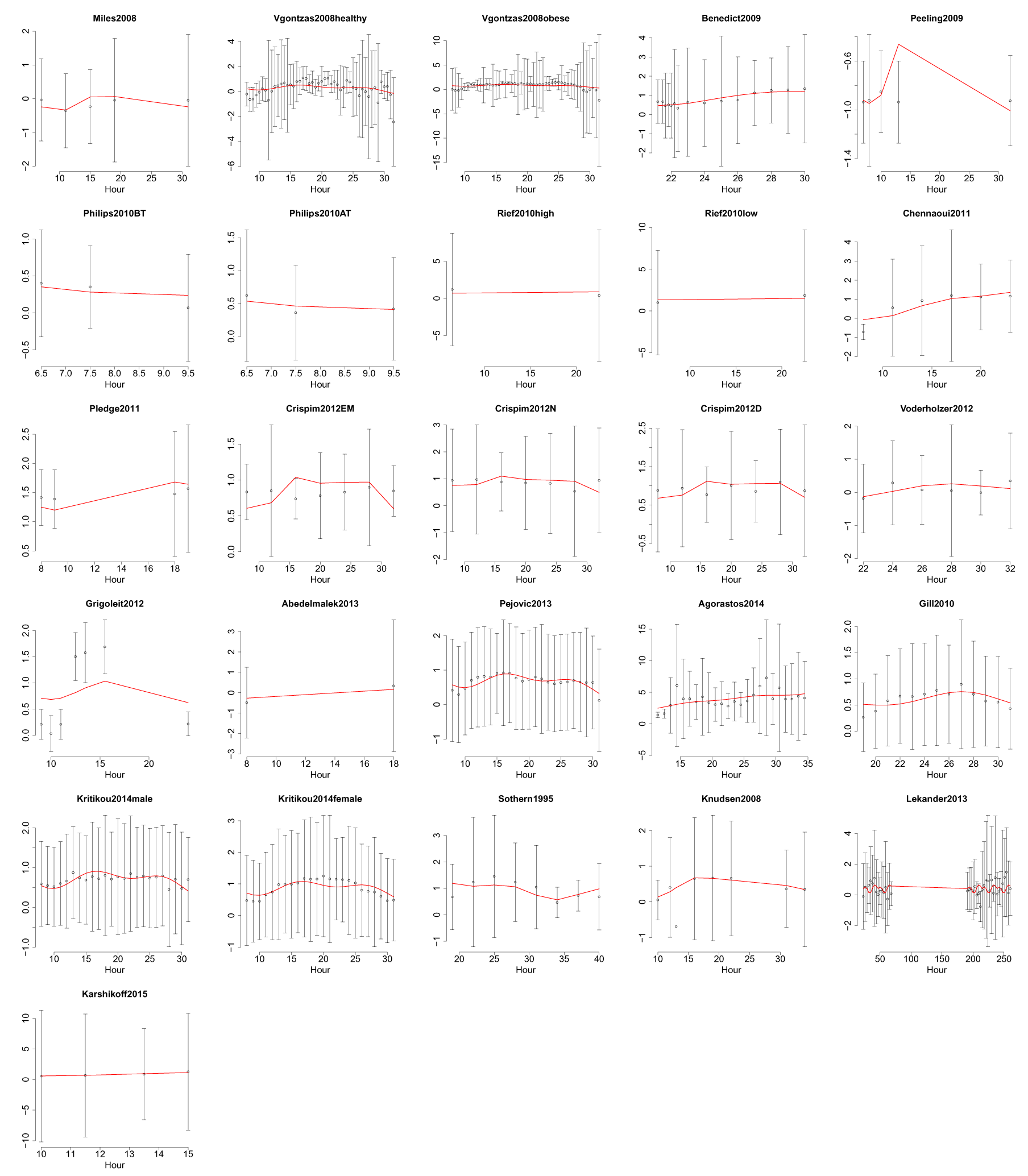
Data and fitted time courses, showing the last 26 out of 56 datasets. Data are shown as estimated from original publications, with error bars showing standard deviations. Y axes show ln IL-6 (pg/ml) throughout. Hours are in chronological time where 1 is 01:00 on the first day. Red lines show predictions from the best-fitting model.

**Fig 7.**
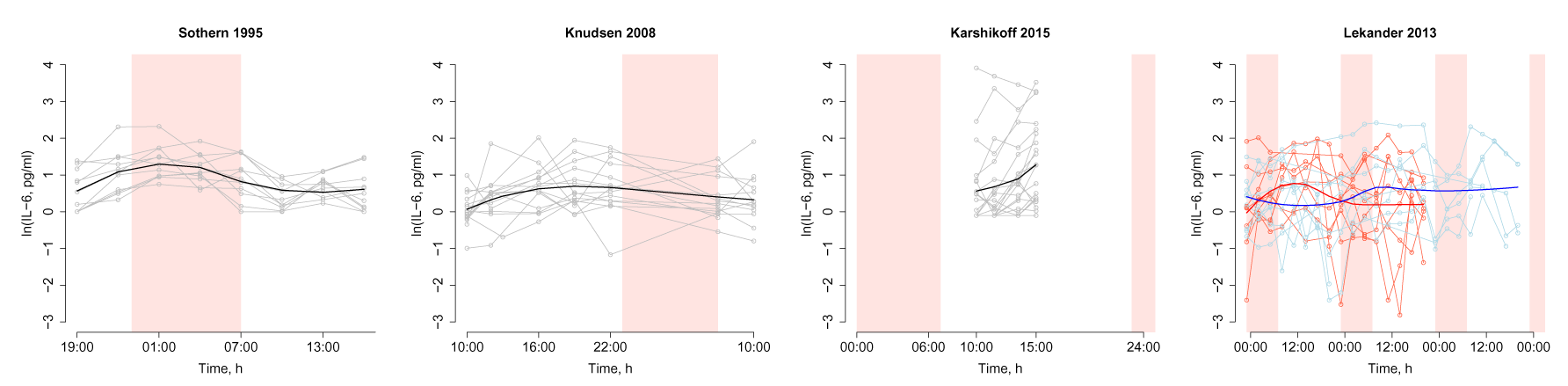
Individual participant data. Individual participant data were available from four datasets and are shown here mainly for the purpose of illustrating the high degree of variability within individuals. To illustrate summary effects within each dataset, thick lines show loess functions fitted to each dataset. Time from 22:00 to 07:00 is shaded to indicate the night. For Lekander 2013, two sets of measurements, made with a few days’ interval, have been plotted over the same time course, in red and blue respectively.

### Assessment of risk of bias

Our literature review identified 36 eligible studies which could not be included. Not counting participants with duplicate data, these studies reported 468 participants, compared to the 1100 included in our meta-analysis. Of the ineligible studies, seven, most of which were published in the 1990:s, were unable to find reliable data because most values were close to or below assay detections limits (Table 2). From the point of view of bias, this is not a major problem since assay detection limits were unrelated to time of day for sampling. More concerning are data that were not reported because no effect was found. Only one paper explicitly stated that data were not shown for this reason [66]. Additionally, an unknown number of studies with eligible data have never been published. It is likely that unreported studies were more likely not to have found significant effects. However, with regard to estimating diurnal phase, we suspect that earlier investigators have been happy to report effects regardless of the location of peaks and troughs, reflected in the wide variety of published estimates (Fig 1). Therefore, even though the first reports of diurnal effects reported peaks in the night, we suspect that the studies included here were not strongly biased towards reporting effects at any particular time of day. Furthermore, to the extent that earlier reports may have been influenced by a prevailing theory, the effect most often referred to is a night-time peak. Since our meta-analysis did not find a night-time peak, we think it is unlikely that a bias in favor of this particular effect will have had a major influence on our model estimates.

Included datasets comprise both data from studies that were designed to measure diurnal or circadian variation, and data from studies that incidentally happened to fulfil our inclusion criteria. The latter group generally had fewer time points for measurements. Since this meta-analysis uses mixed-effects meta-regression by time of day, and is hence not based on single summary measures of included studies, it is not possible to investigate heterogeneity and bias by usual means such as a funnel plot. The results did not strongly depend on any single study. The most influential study was Mehra et al. [44], with a total regression weight of 11.19% over five different datasets.

Since this analysis concerns an observational relationship, confounding from a variety of sources cannot be ruled out. Light exposure and photoperiod were not controlled nor recorded except in laboratory studies, and season or time of the year were not reported frequently enough to justify inclusion in coding. Similarly, physical activity can affect levels of IL-6, but any instructions to participants about physical activity, and measures of their behavior, were for the most part not reported in included studies. Our assumption that participants in datasets not specifying sleep times on average slept 23:00-07:00 would limit the ability to estimate effects of sleep, but that is beyond the scope of this paper, and as the sleep variable was not included in the best-fitting model, the assumption only indirectly affects the risk of bias in diurnal variation. Since the sleep variable made very little difference when added to the best-fitting model, it is unlikely that a different assumption would lead to a different result.

Based on the above considerations, we judge that the risk of bias due to selective publishing and data inclusion is probably moderate to low.

## Discussion

Our meta-analysis confirmed that circulating IL-6 shows diurnal variation. The most marked effect was a morning trough. The best-fitting model included a 24 h and a 12 h cosinor component. The LOESS curve shown in figure 3 suggests a relatively flat curve from the afternoon to the late night, and the better fit obtained by adding the 12 h component may reflect the rather steep change occurring between the morning and the afternoon. For this reason, we are reluctant to consider the best-fitting model as proof that there are two distinct peaks and/or troughs during the 24 h day.

Our analytical approach treats sleep as a confounder to be eliminated, in order to best estimate diurnal variation. The effect of sleep on circulating IL-6 is an interesting question in its own right, but is better addressed by experimental sleep deprivation studies, which were not investigated here. A recent meta-analysis of the effect of sleep deprivation on IL-6 included 12 studies and found no significant effect [20].

One previous meta-analysis [9] (reported again in [10] and [11]) has investigated diurnal variation of circulating IL-6. As discussed in the introduction, this earlier meta-analysis aimed primarily to describe diurnal variation in patients with rheumatoid arthritis, and no systematic method to find data from healthy participants was described. Compared to this earlier meta-analysis, we have used a more systematic approach and we include more data (*k* = 56 datasets compared to k = 11). The present findings contrast markedly with those of the earlier meta-analysis, as that study located a morning peak at approx. 06:00 in healthy controls (Fig 1) as well as in patients with rheumatoid arthritis, while we report a morning trough at about 08:00-09:00 in healthy humans. This raises the hypothesis that diurnal variation is different between healthy humans and patients with rheumatoid arthritis, possibly correlating in patients to the diurnal time course of symptoms such as joint stiffness and pain, which tend to be worse in the morning.

The shape of the diurnal curve estimated in this meta-analysis is not suggestive of a mechanism where the immune system secretes more IL-6 into the blood at night in order to promote sleepiness. While the present results do not disprove this putative mechanism, the lack of a night-time peak, compared to the afternoon, suggests that other regulatory mechanisms are dominant.

The estimated morning trough is rather close after the time of day when cortisol levels peak, and also close to the diurnal trough of monocyte and lymphocyte concentrations in peripheral blood [95–97]. The data investigated here cannot speak directly to the relationship between diurnal variation of IL-6 to cortisol and white blood cell concentrations, and further studies will be required to elucidate whether any direct links exist.

The diurnal variation estimated here is large enough to pose a risk of confounding if sampling is performed without regard to time of day, and we therefore recommend that time of day should be taken in to consideration in studies recording IL-6 in plasma or serum from healthy humans. Further research is required to determine conclusively whether other cytokines also show diurnal variation. As far as we are aware, IL-6 is the the only cytokine to date to be subject to a meta-analysis of diurnal variation.

## Author Contributions

Conceived of the study: GN. Designed the study: GN, MI. Collected data: GN. Analysed data: GN, MI. Interpreted results: GN, TÅ, ML, JA, MI. Drafted the manuscript: GN. All authors read and approved the final version of the manuscript.

